## Supplementary figures and images for "GPRC6A as a novel kokumi receptor responsible for enhanced taste preferences by ornithine"

### Supple. Fig. 2.tif

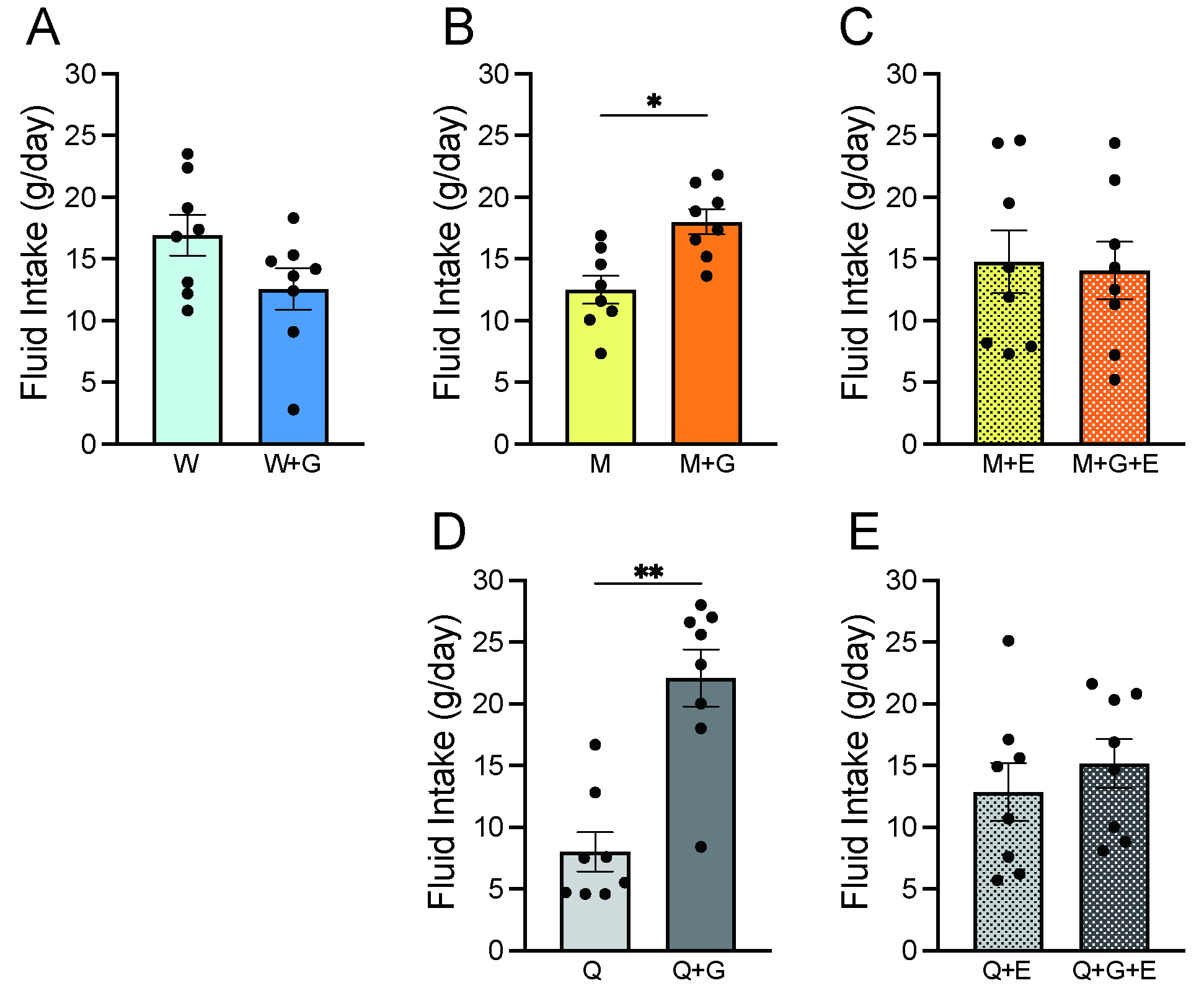

### Supple. Fig. 3-r.tif

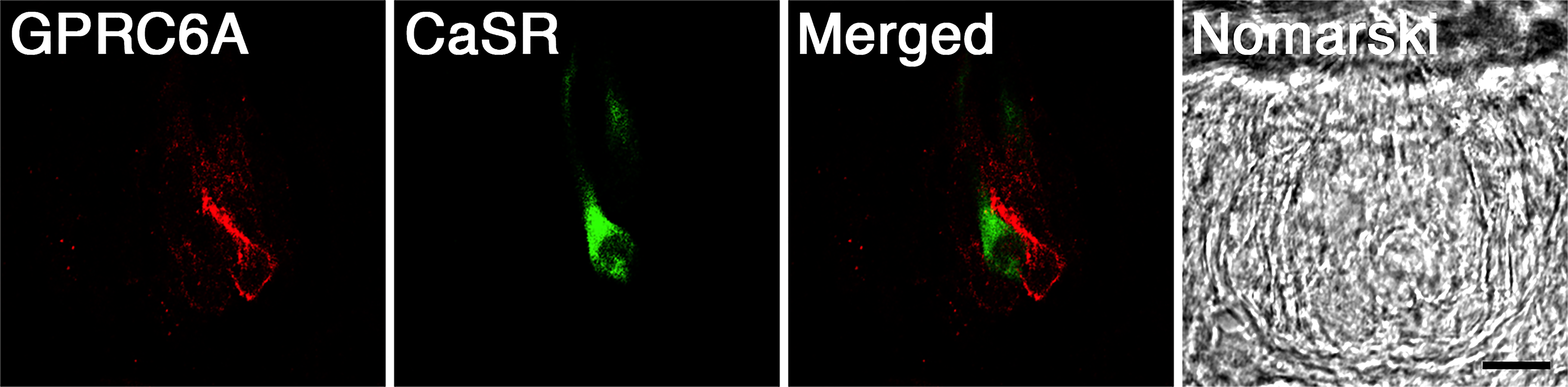

### Supplemental Figure 1

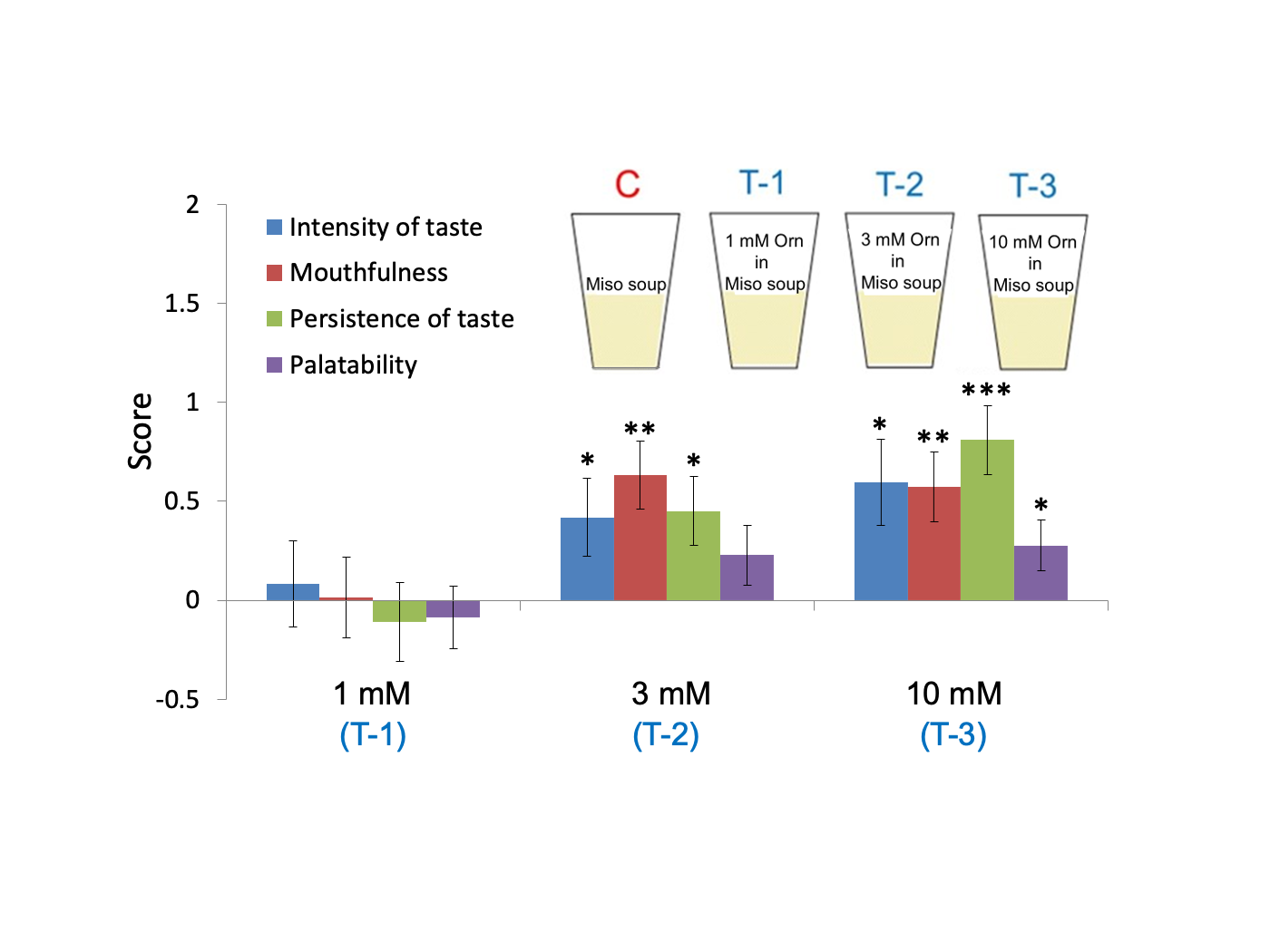
